# Receptor-ligand rebinding kinetics in confinement

**DOI:** 10.1101/370395

**Authors:** A. Erbaş, M. Olvera de la Cruz, J. F. Marko

## Abstract

Rebinding kinetics of molecular ligands plays a critical role in biomachinery, from regulatory networks to protein transcription, and is also a key factor for designing drugs and high-precision biosensors. In this study, we investigate initial release and rebinding of ligands to their binding sites grafted on a planar surface, a situation commonly observed in single molecule experiments and which occurs during exocytosis *in vivo*. Via scaling arguments and molecular dynamic simulations, we analyze the dependence of non-equilibrium rebinding kinetics on two intrinsic length scales: average separation distance between the binding sites and dimensions of diffusion volume (e.g., height of the experimental reservoir in which diffusion takes place or average distance between receptor-bearing surfaces). We obtain time-dependent scaling laws for on rates and for the cumulative number of rebinding events (the time integral of on rates) for various regimes. Our analyses reveal that, for diffusion-limited cases, the on rate decreases via multiple power law regimes prior to the terminal steady-state regime, in which the on rate becomes constant. At intermediate times, at which particle density has not yet become uniform throughout the reservoir, the number of rebindings exhibits a distinct plateau regime due to the three dimensional escape process of ligands from their binding sites. The duration of this regime depends on the average separation distance between binding sites. Following the three-dimensional diffusive escape process, a one-dimensional diffusive regime describes on rates. In the reaction-limited scenario, ligands with higher affinity to their binding sites (e.g., longer residence times) delay the power laws. Our results can be useful for extracting hidden time scales in experiments where kinetic rates for ligand-receptor interactions are measured in microchannels, as well as for cell signaling via diffusing molecules.

## 1 INTRODUCTION

The process of diffusion is a simple way of transporting ligand particles (e.g., proteins, drugs, neurotransmitters, etc.) throughout biological and synthetic media (1, 2). Even though each ligand undergoes simple diffusive motion to target specific or nonspecific binding sites, ensemble kinetics of these particles can exhibit complex behaviors. These behaviors can be traced back to physiochemical conditions, such as distribution of binding sites, concentration of ligands in solution, or heterogeneities in the environment. For instance, biomolecular ligands, such as DNA-binding proteins, can self-regulate their unbinding kinetics via a facilitated dissociation mechanisms dictated by bulk concentration of competing proteins (3–9). Similarly, spatial distribution of binding sites, such as the fractal dimensions of a long DNA molecule (10), or surface density of receptors on cell membranes (11–14) and in flow chambers (15–17), can influence association and dissociation rates of the ligands.

One way of probing these molecular reaction rates is to observe the relaxation of a concentration quench, in which dissociation of ligands from their binding sites into a ligand-free solution is monitored to explore the kinetic rates of corresponding analytes (4, 5, 18–20). Complete time evolution of this relaxation process depends on factors such as chemical affinity between the binding sites and the ligands, dimensions of the diffusion volume, and average distance between the binding sites. While the affinity determines the residence time of the ligand on the binding site (21), the volume available for diffusion can control onset of steady-state regime at which bulk density of the ligands becomes uniform throughout the entire diffusion volume. On the other hand, the spatial distribution of the binding sites can decide how often dissociated ligands revisit binding sites (22). During the nonsteady state at which average concentration of ligands near the binding sites changes with time, these factors can influence the time dependence of the rebinding kinetics in a nontrivial way. In turn, these time dependence can effect the kinetic rates prior to the equilibrium.

This concentration quench scenerio is common both *in vivo* and *in vitro* (Fig. 1). In single-molecule (SM) studies of protein-DNA interactions, short DNA binding sites are sparsely grafted inside a finite-height flow cell (4, 6, 23). The bound proteins may be observed to dissociate into a protein-free solution from their DNA binding sites, allowing measurement of unbinding kinetics. Similarly, in Surface Plasmon Resonance (SPR) apparatus, often more densely packed receptors compared to SM experiments are used to extract kinetic rates. On the other hand, *in vivo* processes, such as exocytosis and paracrine signaling, in which small molecules are discharged into intercellular space to provide chemical communication between cells, can be examples for the relaxation of (effective) concentration quenches (24). Indeed, due to systemic circulation of ligands *in vivo* (e.g., time dependent synthesis/ digestion, or phosphorylation/ diphosphorylation of ligands in cells), a non-steady-state scenario is the dominant situation in biology.

**Figure 1:**
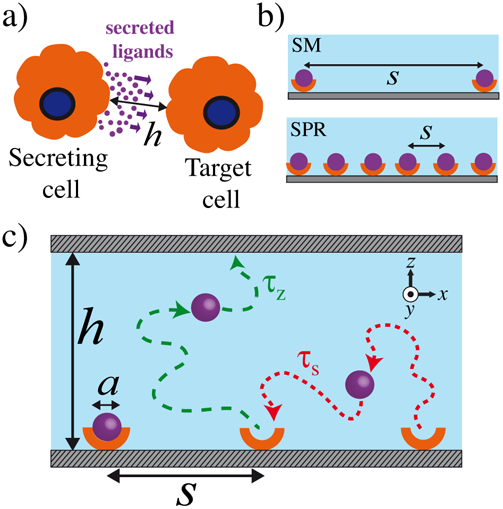
a) Schematics of cell communication via secretion of small ligands into intercellular space of characteristic size of *h*. b) In Single-Molecule (SM) experiments, binding sites (orange) saturated by ligands (purple spheres) are more sparsely distributed compared to SPR experiments. The binding sites are separated by a distance *s*. c) Illustration of diffusion of ligand particles of size *a* initially located at their binding sites. The diffusion volume is confined by two identical surfaces separated by a distance *h*. The particles can diffuse to neighboring binding sites within a diffusion time τ_s_ and to the confining upper surface within a diffusion time of τ_*z*_. Representative trajectories are shown by dashed curves.

Before the steady state is achieved, the separation distance between the binding sites (i.e., grafting density of receptors) influences the kinetic rates by regulating rebinding rates (22, 25). Upon the initial dissociation of a ligand from its binding site into solution, the ligand can return the same binding site (self-binding) or diffuse to neighboring binding sites (cross-binding) (Fig. 1c). In the latter case, the frequency of rebinding events (i.e., on rate for diffusion limited reactions) depends on the average distance traveled by the ligand from one binding site to another. Thus, at time scales comparable to the inter-site diffusion time, the average separation between binding sites becomes a key kinetic parameter.

Experimental studies on ligand-receptor kinetics (26, 27) and signal transduction pathways (13, 28, 29) have highlighted the critical role of the spatial placement of binding sites. In additional studies, in the context of SPR experiments for reservoirs of infinite heights, the effect of correlated rebinding events on the interpretion of dissociation curves has been brought to attention by using a self-consistent mean-field approximation (22, 30). However, in those studies, time scales arising from the diffusion of ligands from neighboring binding sites have not been distinguished due to the one-dimensional nature of the analysis.

Inspired by our own and others’ experiments, as well as the prevalence of the phenomenon in biological systems, our analysis focuses on the time evolution of spontaneous dissociation of an ensemble of Brownian particles from their binding sites into a confined reservoir (Fig. 1c). Using scaling arguments and Molecular Dynamics (MD) simulations, we show that the on rate exhibits two distinct power laws at times longer than initial positional relaxation of the particles but shorter than the time-independent steady-state regime in diffusion-limited reactions. We also derive scaling expressions for the total number of rebinding events experienced by each binding site as a function of time. This quantity can be related to the time-integrated fraction of bound and unbound ligands in experiments (4, 15). Our results indicate that the total number of rebinding events exhibits an unexpected plateau behavior at times much earlier than the onset of the steady state. This plateau regime is terminated by a threshold time scale, which increases with the 4*th* power of the separation distance. Interestingly, this threshold time scale cannot be detected easily in the on rate measurements.

Our scaling expressions were compared to MD simulations of ligands modeled as Brownian particles interacting with their binding sites. A detailed analysis of the simulation trajectories lead to excellent agreements with our scaling predictions. The results that we present here can be applied to new single-molecule studies to determine overlooked time scales involved in binding kinetics and can contribute to our understanding of fundamental principles of the kinetic processes in biological media.

The paper is organized as follows. In Section 2.1, we describe scaling arguments and threshold time scales resulted from our scaling calculations. In Section 2.2, we compare our scaling predictions to MD simulations. We then discuss our results and possible indications for single-molecule studies and biological systems, together with some suggestions for possible experiments in the Discussion section.

## 2 RESULTS

### 2.1 Scaling analysis for ligands diffusing in vertical confinement

Consider *n*_0_ identical particles of size *a* initially (i.e., at *t* = 0) residing on *n*_0_ identical binding sites located on a planar surface at *z* = 0 (Fig. 1c). A second surface at *z* = *h* confines the reservoir in the vertical (i.e., *ẑ*) direction. The size of a binding site is *a*, and the average separation distance between two binding sites is *s*. At *t* = 0, all particles are released and begin to diffuse away from their binding sites into a particle-free reservoir (Fig. 1c). This assumption ignores the finite residence times of ligands on their binding sites and will be discussed further in the following sections.

After the initial release of the ligand particles from their binding sites, each particle revisits its own binding site as well as other binding sites multiple times. The on rate, *k*_on_ (proportional to the local concentration of ligands in diffusion limited reactions), and total number of revisits experienced by each binding site, 𝒩_coll_, reach their equilibrium values once rebinding events become independent of time (i.e., when the ligand concentration in the reservoir becomes uniform). At intermediate times, during which particle concentration in the reservoir is not uniform, various regimes can arise depending on the separation distance *s* or the height of the reservoir *h*.

The time-dependent expressions for *k*_on_ and 𝒩_coll_ prior to the steady state can be related to the length scales of the system on a scaling level after making a set of simplifying assumptions. First, we assume that each particle diffuses with a position independent diffusion coefficient, *D*, without hydrodynamic interactions. We also assume that the particles interact with each other, the binding sites, and surfaces via short-ranged interactions (i.e., interaction range is comparable to the particle size). This approximation is appropriate for physiological salt concentrations, for which electrostatic interactions are short-ranged. We also ignore all prefactors on the order of unity.

After the initial dissociation of the ligand particle, particle can explore a volume *V* (*t*) before it revisits any binding site at time *t*. If there are *ω* binding sites in *V* (*t*), the particle can return any of *ω* binding sites (i.e., *ω* is the degeneracy of the binding sites). Thus, a general scaling ansatz for the on rate can then be written as

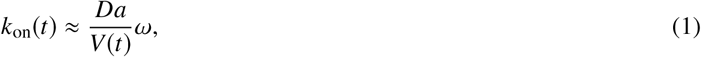

Alternatively, Eq. 1 can also be interpreted as the inverse of the time that is required for a particle to diffuse through *V* (*t*)**/***a*^3^ discrete lattice sites if the diffusion time per lattice site is *D* /*a*^2^. Note that for diffusion-limited reactions *k*_on_(*t*) ~ *c*(*t*), where *c* (*t*) ~ *V* (*t*)^−1^ is the time dependent concentration of ligands. Note that, for simplicity, in Eq. 1, we assume that *ω* has no explicit time dependence although this could be added to the ansatz in Eq. 1. For instance, for binding sites along a fluctuating chain or for diffusing protein rafts on cell membranes, time dependence can be incorporated without losing the generality of Eq. 1.

The total number of rebinding events detected by each binding site at time *t* is related to the on rate as

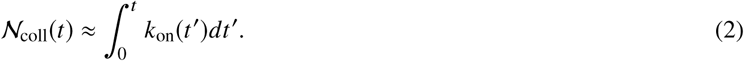

At the initial times of the diffusion of *n*_0_ ligand particles (i.e., *t* ≈ τ_0_ ≈ *a*^2^/ *D*), each particle can undergo a 3d diffusion process to a distance roughly equivalent to its own size (i.e., self-diffusion distance). Since, at 0 < *t* < τ_0_, particles can only collide with their own original binding sites, we have *ω* ≈ 1, and the interaction volume is *V* ≈ *a*^3^ ≈ (τ_0_ *D*)^3/2^. Thus, according to Eq. 1, the on rate is *k*_on_ = 1/τ_0_ and can be considered to be time-independent during *t* < τ_0_ on the scaling level. From Eq. 2, a constant on rate leads to a linearly increasing total number of rebinding events as 𝒩_coll_ ~ *t* (Fig. 2).

**Figure 2:**
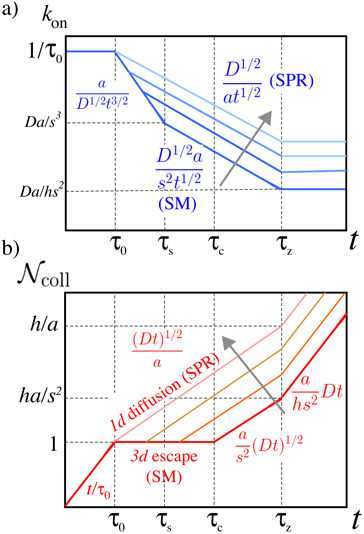
Results of scaling arguments for a) the on rates *k*_on_ and b) the total number of rebinding events 𝒩_coll_ as a function of time in a log-log scale. Arrows indicate the directions of decreasing separation distance between two binding sites (i.e., *s → a*). SM and SPR indicate the regimes related to Single-Molecule and Surface Plasmon Resonance experiments, respectively. The threshold time scales refer to onset for darker lines. See also Table 1 for the definition of the threshold times.

For *t* > τ_0_, each particle can diffuse to a distance *r* > *a*. If the separation distance between the binding sites is *s* ≈ *a*, particles can visit any of the nearest binding sites at *t* ≈ τ_0_. If the separation distance is large (i.e., *s* ≫ *a*), particles can travel to neighboring sites only after a time τ_s_ ≈ *s*^2^/*D*, at which the average distance traveled by any particle is *s*. At τ_0_ < *t* < τ_s_, individual particles perform 3d diffusion, and thus, the interaction volume is given by *V* (*t*) ≈ (*Dt*)^3/2^. Since the volume experienced by particles is *V* (*t*) < *s*^3^ at *t* < τ_s_, on average one binding site is available per particle in the interaction volume, thus, *ω* ≈ 1. Thus, using Eq. 1, we can obtain *k*_on_ ~ *t*^−3/2^.

Interestingly, at τ_0_ < *t* < τ_s_, the number of revisits per binding site, 𝒩_coll_, does not increase since most particles are on average far away from their own and other binding sites. On the scaling level, this results in a plateau behavior for the cumulative collision number (i.e., 𝒩_coll_ ≈ 1), as illustrated in Fig. 2. Note that plugging *k*_on_ ~ *t*^−3/2^ into Eq. 2 leads to a weak explicit time dependence for the 𝒩_coll_ at 0 < *t* < τ_s_ (i.e., 𝒩_coll_ ~ 1 + *t*^−1/2^). The *t*^−1/2^ dependence indicates that 𝒩_coll_ stays almost constant during this regime.

At *t* > τ_s_, the particles can encounter other neighboring binding sites apart from their own, thus, *ω* > 1. The particle density near the bottom surface of the reservoir is nearly uniform, but the overall density is still non-uniform throughout the reservoir. This can be seen in the simulation snapshots shown in Fig. 3 (we will discuss our simulation results further in the next section). Only at a threshold time dictated by the height of the reservoir, τ_*z*_ ≈ *h*^2^/*D*, each particle on average reaches the physical limits of the reservoir, and on rate does reach its steady-state limit (i.e., *k*_on_ ≈ *Da*/*hs*^2^), as shown in Fig. 2. At earlier times, *t* < τ_*z*_, since there are ligand-free regions in the reservoir (Fig. 3), *V* (*t*) and thus, on rate still must exhibit a time dependence.

**Figure 3:**
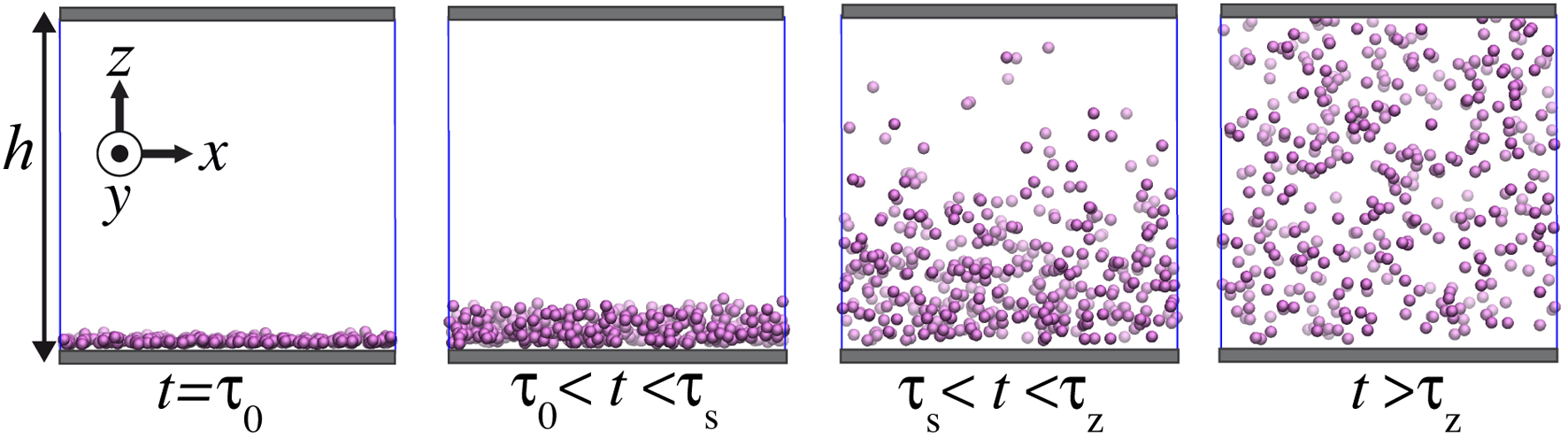
Simulation snapshots at various time windows showing the time evolution of the particle concentration throughout a simulation box of height *h*/*a* = 50. The separation distance between the binding sites is *s*/*a* = 2.5. The blue lines indicate the borders of the original simulation box. Periodic boundary conditions are applied only in the *x*̂ and *y*̂ directions, whereas the reservoir is confined in the *z*̂ direction by two identical surfaces.

One way of obtaining the time-dependent on rate at τ_s_ < *t* < τ_*z*_ is to consider the diffusion of a single-particle and use *V* (*t*) *≈* (*Dt*)^3/2^ and *ω* ≈ (*Dt* /*s*^2^) for the number of binding sites per an area of (*Dt*). Consequently, Eq. 1 leads to *k*_on_ ≈ *D*^1/2^ *a*/*s*^2^*t*^1/2^ ~ *t*^−1/2^. This scaling is due to quasi one-dimensional propagation of the particle cloud across the reservoir although each particle undergoes a 3d diffusion process (Fig. 3). Alternatively, to obtain the scaling form for on rate, one can consider the overall diffusion of the particle cloud at *t* > τ_s_ (Fig. 3). Then, the total explored volume scales as *Vt*) ~ (*Dt*)^1/2^, and the total number of binding sites in this volume is *ω* ~ 1/*s*^2^. Thus, Eq.1 leads to *k*_on_ ~ *t*^−1/2^.

The rapid drop of on rate at *t* > τ_0_ with multiple negative exponents has consequences on 𝒩_coll_. According to Eq. 2, the scaling form for 𝒩_coll_ at *t* > τ_s_ can be obtained from the partial integration of the corresponding on rate expressions for appropriate intervals (Fig. 2a). Thus, at *t* > τ_s_, the total number of rebinding, on the scaling level, is

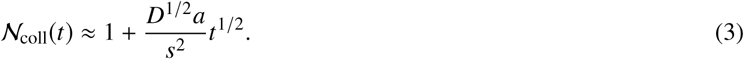

We note that inserting *t* = τ_s_ into Eq. 3 leads to 𝒩_coll_ ≈ 1 for any value of *s*/*a* > 1 since the second term on the right hand side of Eq. 3 is smaller than unity. This indicates that the plateau regime predicted for 𝒩_coll_ at *t* < τ_s_ persists even at *t* > τ_s_ (Fig. 2b). Only at a later threshold time, τ_c_ > τ_s_, the second term of Eq. 3 becomes considerably larger than unity, and so does the number of revisits, 𝒩_coll_. The threshold time, τ_c_, can be obtained by applying this result on the second term of Eq. 3 (i.e.,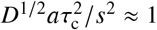), which provides an expression for the terminal time of the plateau regime as

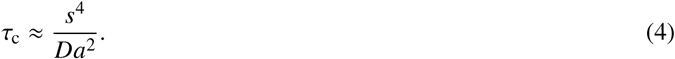

At *t* > τ_c_, the total number of revisits per binding site begins to increase above unity. The functional form of this increase at τ_c_ < *t* < τ_*z*_ can be obtained by integrating the on rate (i.e., *k*_on_ ~ *t*^−1/2^) as 𝒩_coll_ ~ *t*^1/2^. This sublinear increase of 𝒩_coll_ continues until the particle density becomes uniform throughout the entire reservoir at *t* = τ_*z*_. At later times *t* > τ_*z*_, diffusion process obeys Einstein-Smoluchowski kinetics, where the on rate reaches its time-independent steady-state value, and where 𝒩_coll_ increases linearly (Fig. 2).

To summarize, as also schematically illustrated in Fig. 2, according to our scaling analysis, at *t* < τ_0_, the on rate is constant due to self-collisions with the original binding site. At later times, the on rate decreases as *k*_on_ ~ *t*^−3/2^ until *t* < τ_s_ due to the 3d escape process of particles away from their binding sites (Fig. 2a). Once particles diffuse to distances on the order of *s*, the particle cloud diffuses in a 1d manner, and the on rate decays with a slower exponent, *k*_on_ ~ *t*^−1/2^. When the particles fill the reservoir uniformly, a steady-state value of *k*_on_ ~ *a*/(*hs*^2^) takes over. Interestingly, at the threshold time τ_c_, at which we predict a crossover for 𝒩_coll_, the on rate does not exhibit any alterations and continues to scale as *k*_on_ ~ *t*^−1/2^.

The regime during which 𝒩_coll_ is independent of time on the scaling level is smeared out in the limit of *s → a* as shown in Fig. 2b. If *s* = *a*, the plateau in 𝒩_coll_ completely disappears, and a scaling 𝒩_coll_ ~ *t*^1/2^ determines the cumulative rebinding events at τ_0_ < *t* < τ_*z*_. This indicates that the 3d escape process disappears, and a 1d diffusion-like behavior prevails after the initial dissociation of ligands. This behavior is common in SPR experiments, where receptors are often densely grafted.

In the equations below, the scaling expressions for the on rates rescaled by 1/τ_0_ ≈ (*D*/*a*^2^)^−1^ and the total number of revisits are given together with their respective prefactors for corresponding time intervals (see Table 1) as

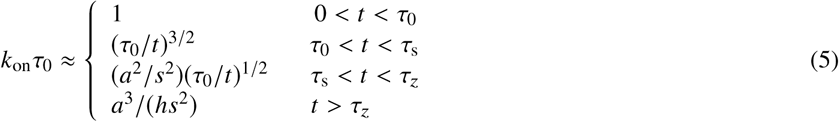

**Table 1:** The threshold times and their scaling expressions with numerical estimates for various systems. In the estimates, a ligand of size *a* = 1 nm and a diffusion coefficient of *D* = 100 *μ*m^2^/s are assumed. The estimates are for (1) *s* = 10 nm and *h* = 10^2^ *μ*m (e.g., SPR case), (2) *s* = 1 *μ*m and *h* = 10^4^ *μ*m (e.g., SM case (4)), (3) diffusion of insulin secreted from isolated vesicles into intercellular space of height *h* = 30 nm and *s* = 10 nm (determined from the avarege insulin concentration of ≈ 40 mM in the vesicle (31)), (4) release of ≈ 10 *μ*M of GTP*γ*S (i.e., *s* = 100 nm) from a eosinophils-cell vesicle (32) with an average cell-to-cell distance of *h* = 100 μm (i.e., ca. 500 cells per microliter).

|  | Scaling | SPR<sup>(1)</sup> | SM<sup>(2)</sup> | Exocytosis<sup>(3)</sup> | Exocytosis<sup>(4)</sup> |
| --- | --- | --- | --- | --- | --- |
| $\tau_0$ | $a^2/D$ | $10^{-9}$ s | $10^{-9}$ s | $10^{-9}$ s | $10^{-9}$ s |
| $\tau_s$ | $s^2/D$ | $10^{-6}$ s | $10^{-2}$ s | $10^{-6}$ s | $10^{-4}$ s |
| $\tau_c$ | $s^4/a^2D$ | $10^{-4}$ s | $10^4$ s | $10^{-4}$ s | $10^0$ s |
| $\tau_z$ | $h^2/D$ | $10^2$ s | $10^6$ s | $10^{-5}$ s | $10^2$ s |

Similarly, for the total number of revisits

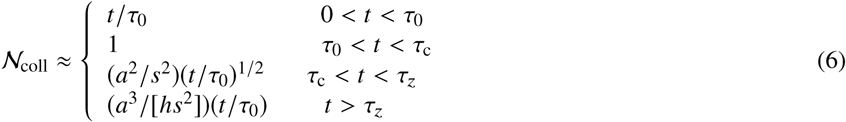

In Table 1, we provide some numerical examples for the above times scales approximately corresponding to SPR and SM experiments, and exocytosis. Note that the scaling expression given in Eqs. 5 and 6 can also be obtained by considering the relaxation of a Gaussian particle distribution in corresponding dimensions (see Appendix section).

In the next section, we will compare our scaling arguments with the coarse-grained MD simulations and investigate the relation between threshold time scales, and the two length scales *s* and *h*.

### 2.2 Comparison with molecular dynamics simulations

#### 2.2.1 Description of simulation methodology

Coarse-grained MD simulations were performed by using the LAMMPS package (33). The ligands that are modeled by spherical beads of size *a* were placed on a rigid surface with a prescribed seperation distance of *s*. The surface is also composed of the same type of beads as ligands. The ligands interact with each other and the surfaces via a short range Lennard-Jones potential (see Eq. 14 in Appendix section) in an implicit background solvent at constant temperature (34). A prescribed number of ligands (i.e., *n*_0_ = 400 – 6400) are allowed to diffuse into a confined reservoir at *t* > 0. The reservoir is periodic in the lateral directions but confined in the vertical direction by a second surface identical to the first one (Fig. 3).

The rescaled height of the simulation box *h*/*a* and the rescaled separation distance *s*/*a* were separately varied to monitor their effects on time dependencies of *k*_on_(*t*) and 𝒩_coll_ (*t*). In the extraction of *k*_on_ values, the binding sites are defined as the initial positions of ligands at *t* = 0. Any particle that is found within the collision range of any binding site (i.e., *r_c_*/*a* = 2^1/6^) at a given time *t* is counted as a bound particle. In our analyses, *k*_on_ (*t*) is defined as the normalized fraction of binding sites occupied by ligands for diffusion-limited reactions. For reaction limited case, *k*_on_ (*t*) corresponds to raw dissociation data. The values of 𝒩_coll_ (*t*) were calculated via Eq. 2. All simulations were carried out until the calculated on rates reached their respective steady states (see Appendix section for further simulation details).

#### 2.2.2 Diffusion-limited kinetics

We first consider the scenario for which the reactions between the binding sites and ligands are diffusion limited. Thus, the average residence time of the ligand on the binding site is on the order of τ_0_, which is the self-diffusion time of a particle in the simulations. We achieved this by using a purely repulsive WCA potential (35) with a cut-off distance of *r_c_*/*a* = 2^1/6^ (Eq. 14 in the Appendix section). This setup, as we will see, allows us to observe the regimes predicted in Section 2.1 more clearly. We will further discuss the longer residence times in conjunction with other time scales in the following sections.

##### Qualitative analyses of simulations

In Fig. 3, we present a series of simulation snapshots to demonstrate the diffusion process of *n*_0_ = 400 particles over the time course of the simulations, for *h*/*a* = 50 and *s*/*a* = 2.5. These numbers lead to characteristic times ranging from τ_s_ ≈ τ_0_ to beyond τ_*z*_ ≈ 10^4^τ_0_ for the system shown in Fig. 3. At short times, *t ≈* τ_0_, the particles are mostly near the reactive (bottom) surface as can be seen in Fig. 3. As the time progresses, the particle cloud diffuses vertically to fill the empty sections of the box. At *t* < τ_*z*_, the particle density near the surface changes with time, and visually the concentration is not uniform in the box. Only for *t* > τ_*z*_, the particle density becomes uniform, and the initial concentration quench is completely relaxed as illustrated in Fig. 3.

##### Densely placed binding sites in finite-height reservoirs

To systematically compare our scaling predictions with the simulations, we fixed the separation distance to *s*/*a* = 2.5 and varied the height of the reservoir. Fig. 4 shows the data calculated from the particle trajectories for the rescaled on rate *k*_on_τ_0_ and the total number of revisits 𝒩_coll_ as a function of the rescaled simulation time *t*/τ_0_. At short times (i.e., *t* < τ_0_), during which particles can diffuse only to a distance of their own size, 𝒩_coll_ increases linearly, whereas on rates *k*_on_ have no or weak time dependence in accord with our scaling calculations. In Fig. 4a, at approximately *t* ≈ τ_0_, we observe a rapid drop in *k*_on_, which is described nominally by an exponential function [exp(–*t*/τ_0_)] (dashed curve in Fig.4a). However, we should also note that, in the system presented in Fig. 4a, *s*/*a* = 2.5, thus, τ_s_ ≈ 6τ_0_. Hence, the decay in the on rate is arguably the beginning of a power law with an exponent approximately 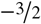 (Fig. 2a).

**Figure 4:**
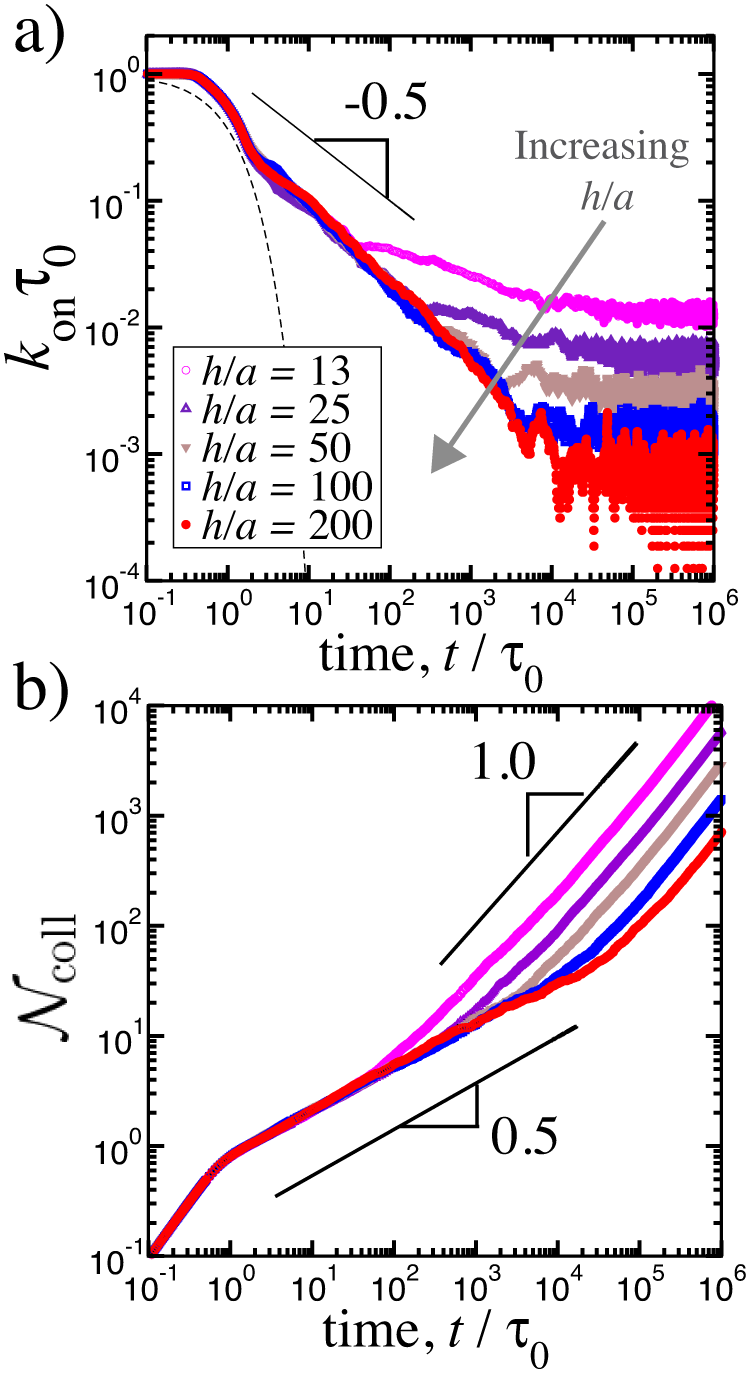
a) Rescaled on rates as a function of the rescaled simulation time for various reservoir heights. The distance between two binding sites is *s*/*a* = 2.5. A running average over 20 data points is shown for all cases for clarity. b) Total number of revisits per binding site obtained via Eq. 2 by using non-averaged data series of (a). For all cases, number of ligands particles is *n*_0_ = 400.

According to our scaling analysis, for small enough separation distances (i.e., *s* ≈ *a*), the on rate obeys a power law (i.e., *k*_on_ ~ *t*^−1/2^) at the intermediate times, τ_s_ < *t* < τ_*z*_ (see Fig. 2a). In Fig. 4a, a slope of –0.56 ± 0.04 describes the decay of the on rates in accord with our scaling prediction. We have also tested larger systems with *n*_0_ = 1600 and *n*_0_ = 6400 particles and obtain similar exponents. At longer times (i.e., *t* > τ_*z*_), the on rates in Fig. 4a reach their respective steady-state values at *t* = τ_*z*_ = *h*^2^/*D*, which depends on the equilibrium concentration of the ligands (i.e., *k*_on_ ~ 1/*hs*^2^). That is, for a fixed *s*/*a*, increasing the height *h*/*a*, decreases the concentration and thus the steady-state values of the on rates as seen in Fig. 4a.

As for the total number of revisits, 𝒩_coll_, in Fig. 4b, the simulation results for densely-packed binding sites show a power law dependence on time as 𝒩_coll_ ~ *t*^0.44±0.05^ at the intermediate times in accord with the prediction (Fig. 2b). Once this regime ends, a subsequent terminal linear regime, in which 𝒩_coll_ ~ *t*^1.0^, manifests itself in Fig. 4b. According to our scaling analysis (Eq. 6), the onset of this long-time linear regime is set by τ_*z*_. Thus, increasing the height of the reservoir *h*, only shifts the onset to later times (Fig. 4b). Note that in Fig. 4b, for small values of *h*/*a*, the exponent is more close to unity since it takes less time to reach a uniform ligand density in smaller simulation boxes.

##### Effect of separation distance on rebinding kinetics

As discussed in Section 2.1, prior to the steady state, diffusion time between binding sites significantly affects the apparent dissociation kinetics of ligands. To study this phenomenon, we ran simulations with various values of the separation distance ranging from *s*/*a* = 2.5 to *s*/*a* = 50 for a fixed height of *h*/*a* = 50 (Fig. 5). While the short-time kinetic behaviors in Fig. 5 are similar to those in Fig. 4 regardless of the surface separation, the long-time behavior exhibits various regimes depending on the separation distance *s*/*a* in the simulations.

**Figure 5:**
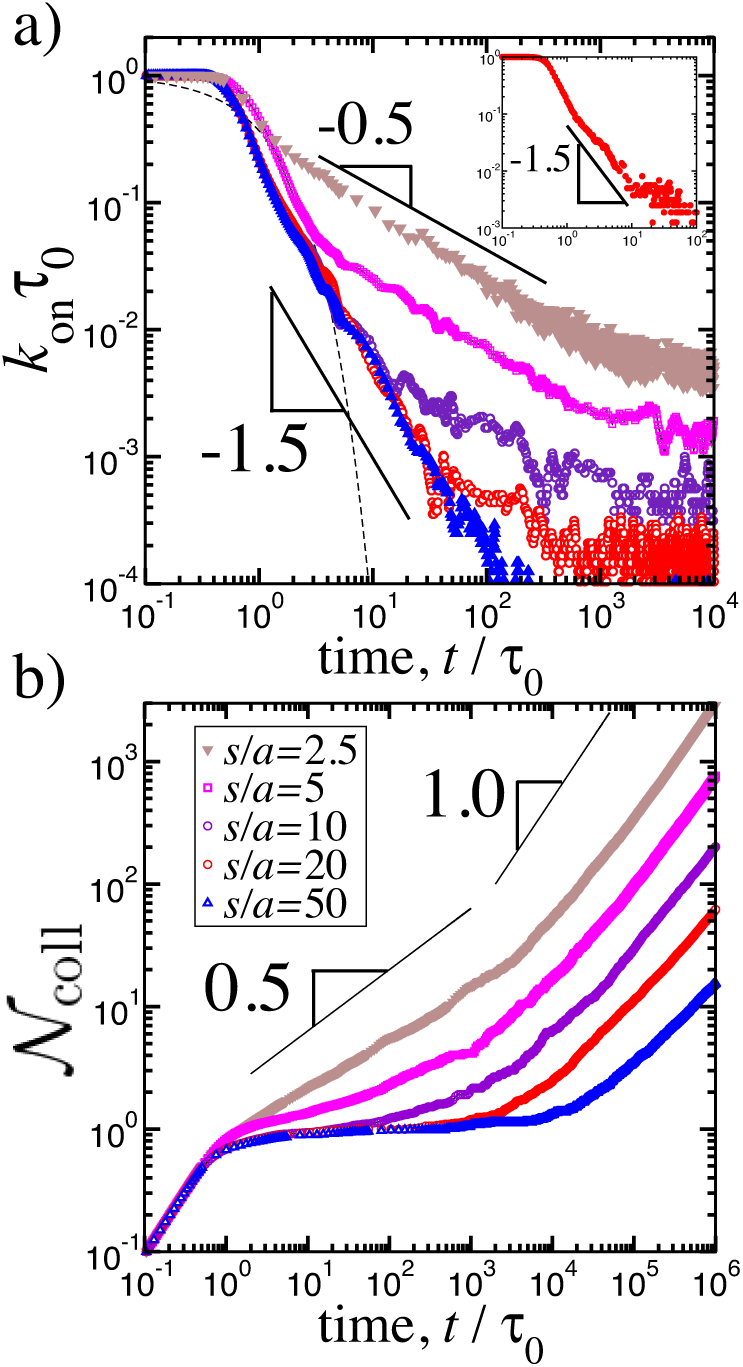
a) On rates rescaled by the unit diffusion time τ_0_ as a function of rescaled time for various separation distances between binding sites for a fixed box height of *h*/*a* = 50. Each data set is averaged over 3 to 5 separate simulations of the systems containing *n*_0_ = 400 – 6400 particles. For clarity, running averages over 10 points are shown. Inset shows the on rate for a system composed of *n*_0_ = 1600 particles for *s*/*a* = 10 and *h*/*a* = 1000 b) The total number of rebinding events obtain via Eq. 2 for *n*_0_ = 400-particle systems by using non-averaged data sets.

For *s*/*a* ≈ 1, as discussed earlier, a slope close to −½ can describe the decay of the on rates before the time-independent steady state (Fig. 5a). For *s*/*a* ≫ 1, this slope is replaced by a stronger decay *k*_on_ ~ *t*^−1.46±0.13^ at *t* > τ_0_ (Fig. 5a). In our scaling analysis, the exponent 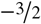 controls the decay of on rate until the threshold time scale of τ_s_ sets in (Fig. 2a). Indeed, in Fig. 5a, as *s*/*a* is increased, the scaling *k*_on_ ~ *t*^−1.5^ replaces *k*_on_ ~ *t*^−0.5^ type of behavior gradually. For the intermediate values of *s*/*a* (i.e., *s*/*a* = 5, 10, 20, 50 in Fig. 5a), this transition can be observed in accord with the scaling prediction in Fig. 2a. In the inset of Fig. 5a, we also show a system with *s*/*a* = 10 and *h*/*a* = 1000 for a larger system of *n*_0_ = 1600 particles; The 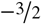 exponent is more apparent since the two threshold times, τ_*z*_ and τ_s_, are well separated due to the large ratio of *h*/*s* ≫ 1.

##### Emergence of plateau behavior in total rebinding events for sparsely placed binding sites

The data in Fig. 5b shows the distinct behavior of 𝒩_coll_ for *s*/*a* ≫1 as compared to the cases, where binding sites are closer to each other (Fig 4b). As discussed earlier in Fig. 4b, for *s*/*a* ≈ 1, a slope around 𝒩_coll_ ∼ *t*^0.44^ is dominant at *t* < τ_*z*_. However, for *s*/*a* ≫ 1, a plateau regime replaces this behavior at the intermediate times (Fig. 5b). As *s*/*a* is increased, the plateau regime becomes broader by expanding to longer times. This trend is also in agreement with our scaling analyses (Fig. 2b).

The plateau regime in 𝒩_coll_ is followed by an incremental behavior as seen in Fig. 5b. The predicted power law following the plateau is 𝒩_coll_ ~ *t*^1/2^ for τ_s_ < *t* < τ_*z*_ (Fig. 2b). Within the duration of our simulations, we observe a mixture of slopes instead of a single exponent of ½. For instance, for *h*/*a* = 50, the slope that we can extract at long times is smaller than unity but larger than ½ since τ_s_ ≈ τ_*z*_ (blue triangles in Fig. 5b). This is due to the small ratio of the two threshold time scales, τ_*z*_ /τ_c_ = (*ha*/*s*^2^)^2^ ≈ 10, for the data shown in Fig. 5b.

To further separate these two time scales, we performed simulations for a fixed *s*/*a* = 10 and for various values of *h*/*a* = 50 − 2000. The results are shown in Fig. 6 for *t* > τ_s_ ≈ 100τ_0_. For all the data sets in Fig. 6, τ_s_ and τ_c_ are identical (i.e., equal *s*/*a*). Thus, only difference in their kinetic behavior arises due to the variations in *h*/*a*, which in turn determines the duration of the τ_*z*_ – τ_c_ interval. In Fig. 6, ideally, the regime with 𝒩_coll_ *t*^1/2^ should be observable at τ_s_ < *t* < τ_*z*_. However, we rather observe a weaker increase before a slope of around ½ emerges. We attribute this behavior to the inherent weakness of scaling analyses since even at *t* > τ_s_, ligands can collide with multiple binding sites frequently enough, particularly for small separation distances. These collisions, in turn, can result in a slight increase in 𝒩_coll_ similar to that observed in Fig. 6.

**Figure 6:**
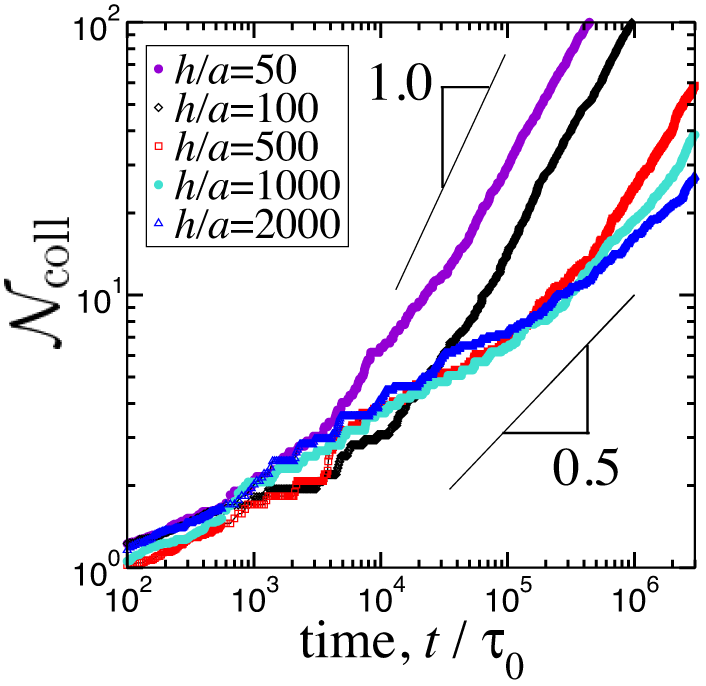
The total number of rebinding event as a function of rescaled simulation time for various rescaled heights at a fixed separation distance *s*/*a* = 10. For clarity, only the data for *t* > τ_s_ ≈ 100τ_0_ is shown. For all cases, number of ligands is *n*_0_ = 400.

At long times *t* > τ_*z*_, 𝒩_coll_ increases with an exponent around unity in the simulations (Figs. 5b and 6) in accord with a time-independent *k*_on_. Note that, for simulations longer than performed here, which are not feasible for computational reasons, we anticipate a convergence to a slope of unity for all of our configurations.

##### Threshold time for 𝒩_coll_ plateau

We also performed a separate set of simulations to specifically identify the scaling dependence of τ_c_ on the separation distance (Eq. 4). We fix the ratio *h*/*s* = 10 and vary the separation distance between *s*/*a* = 10 – 100 and the height between *h*/*a* = 100 – 1000. We fit the data encompassing the time interval τ_0_ < *t* < τ_*z*_ to a function in the form of *f*(*t*)= 1 + (*t*/τ_s_)^1/2^ to extract the threshold time τ_c_ at which plateau regimes ends. The results shown in Fig. 7 are in close agreement with our scaling prediction; The data can be described by a scaling τ_c_ ~ *s*^3.5±0.5^. The fact that the exponent extracted from the simulations is smaller than 4 but larger than 2 in Fig. 7 indicates that the terminal threshold time for the plateau, τ_c_, is distinct and well-separated from τ_s_.

**Figure 7:**
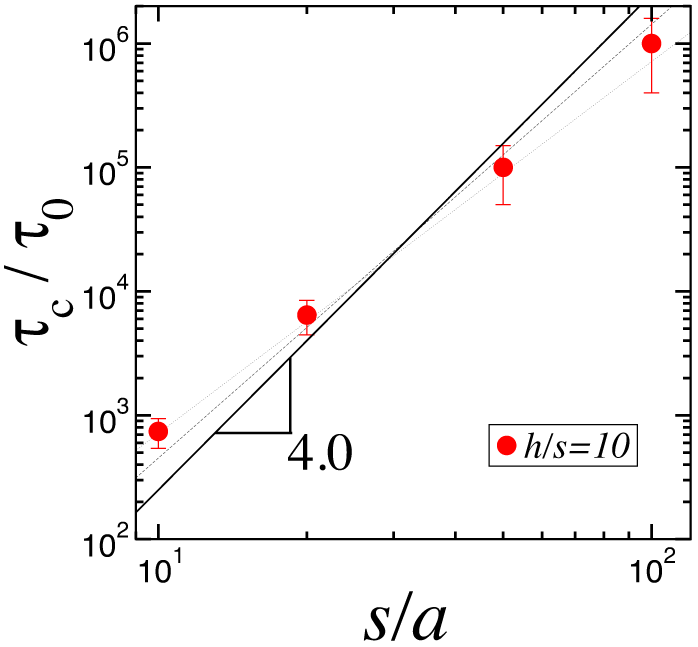
Log-log plot of τ_c_ extracted from simulations by fitting the respective intervals to a function in the form of *f* (*t*) = 1 + (*t*/τ_c_)^1/2^ for *h*/*s* = 10 as a function of the rescaled separation distance. The thin lines show the slopes of 3.0 and 3.5. For the data sets, *h*/*a* = 100, 200, 500, 1000, and for *s*/*a* = 10, 20, 50, 100. For all cases, the number of ligands is *n*_0_ = 400.

#### 2.2.3 Effect of Reaction-limited kinetics

In Section 2.2.2, we consider the diffusion limited case, where being within the collision range of a binding site is enough to be counted as bound for any ligand. That is τ_off_ ≈ τ_0_. However, most molecular ligands including DNA binding proteins can have finite lifetimes on the order of minutes to hours (4, 5). Long residence times can indeed intervene with the threshold times and regimes predicted by our scaling arguments.

To test how the finite residence times can effect the rebinding rates, we ran a separate set of simulations, in which a net attraction is introduced between the binding sites and the ligands, for two different separation distances, *s*/*a* = 2.5 and *s*/*a* = 20, with *h*/*a* = 50 (Fig. 8). The attraction was provided by increasing the cutoff distance and varying the strength of the interaction potential in the simulations (see Appendix for details). As a result of this net attraction, the ligands stay on their binding sites for longer times (i.e., τ_off_ > τ_0_). Importantly, the data presented in Fig. 8 corresponds to the fractions of occupied binding sites since on rate is no more proportional to the concentration in the reaction-limited case.

**Figure 8:**
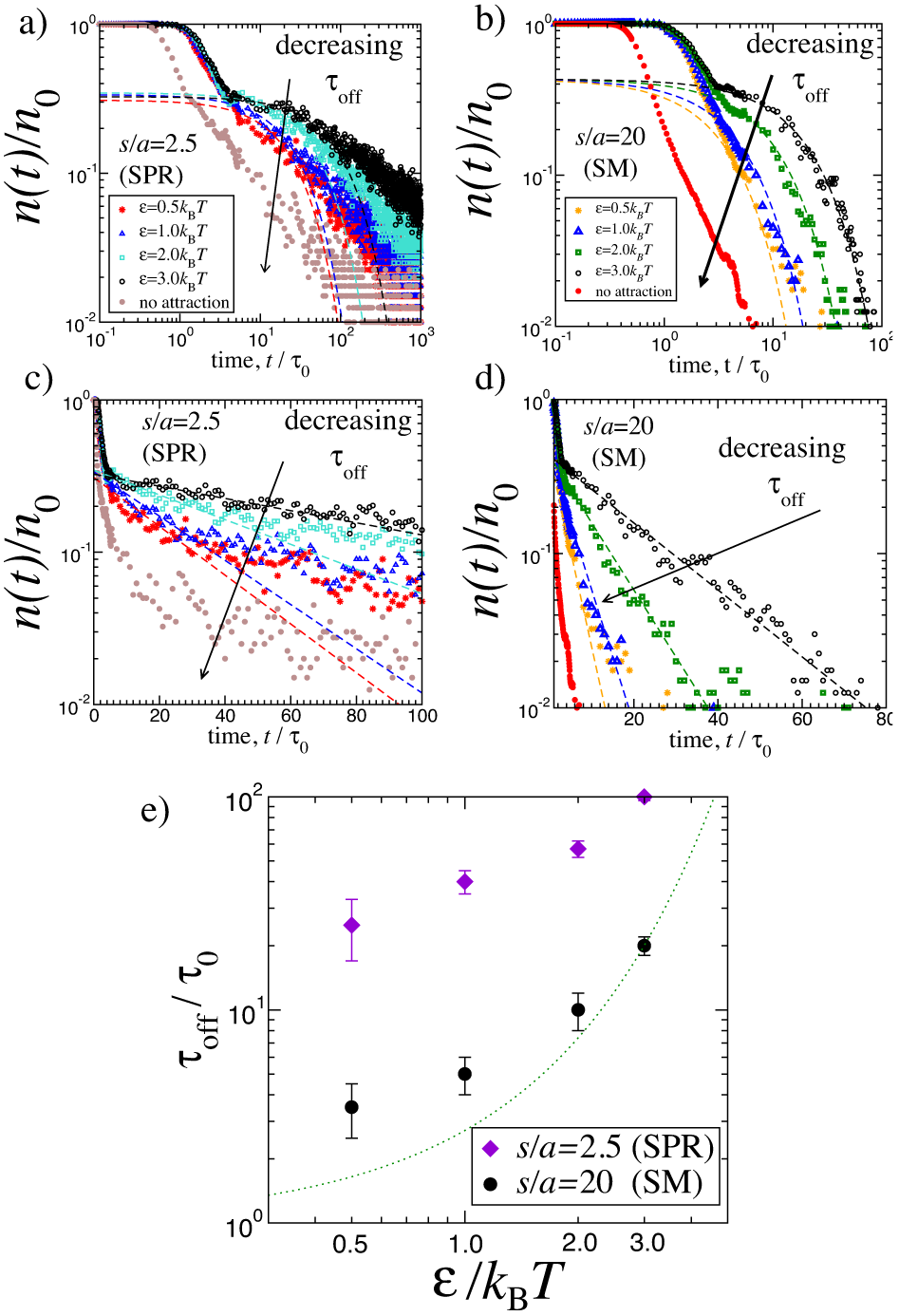
Fraction of binding sites occupied by ligands obtained from the simulations in which a net attraction between binding sites and ligand particles leads to finite residence times prior to dissociation. The strength of attractions is ∊ (in the units of *k*_B_*T*) (see Appendix for the pair potential). The brown and red data sets are the same as those in Figs. 4 and 5 with no net attraction. a) *s*/*a* = 2.5 and b) *s*/*a* = 20, for *h*/*a* = 50. c) and d) log-linear plot of the data sets in (a) and (b). The dashed curves are exponential fits (≈ exp(−*t*/τ_off_)) to the data sets for *t* ≳ 3τ_0_. e) The log-log plot of residence times obtained from plots a-d as a function of the attraction strength of the interaction potential for *s*/*a* = 2.5 and *s*/*a* = 20. The dotted curve is *f* (*x*) = exp(*x*/*k*_B_*T*).

In Fig. 8, at the short times, we observe a rapid drop, regardless of the strength of the attraction. We attribute this common initial behavior to the escape process of the ligands from the attractive potential. After the rapid decay, for high affinities (longer lifetimes), the regimes with either –½ or 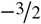 exponents disappear and an exponential function in the form of exp (–*t*/ τ_off_) can describe the data at the intermediate times (dashed curves in Fig. 8a-b). This can also be seen in the log-linear plots in Fig. 8c and d. As the attraction strength is decreased, the power laws become dominant again as expected from the diffusion limited cases (Fig. 8a-d). For longer simulations, we anticipate that the power laws should be attainable if *s* and *h* are large enough. This can be seen in Fig. 8a; after the exponential decay at around *t* = 100τ_0_, a slope of –½ begins to emerge. We will further discuss the criterion for observing an exponential decay in the Discussion section (Fig. 9)

**Figure 9:**
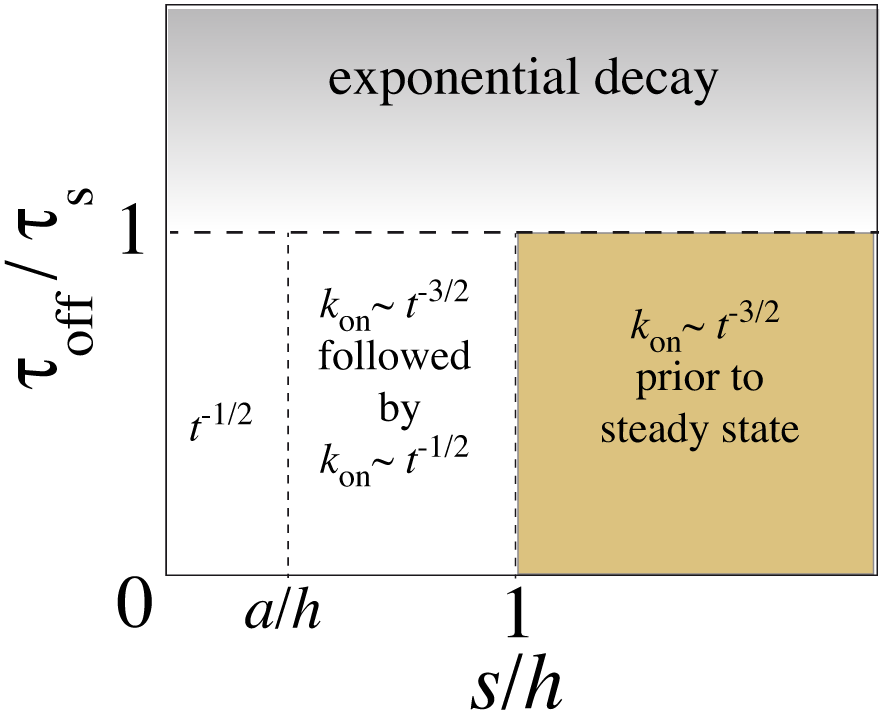
Summary of on rate scaling behaviors predicted for various systems. Along the dashed lines, on rate follows a −½ exponent. The residence time of a ligand on its binding site is defined as the inverse off rate, τ_off_ ≡ 1/*k*_off_.

In Fig. 8e, the residence times τ_off_ extracted from the exponential fits are shown for two separation distances, *s*/*a* = 2.5 and *s*/*a* = 20. Even though the attraction strengths between the ligands and binding sites are identical for two cases (i.e., *ε* = 3, 2, 1, 0.5*k*_B_*T*), the extracted lifetimes are longer for the smaller separation distance (Fig. 8). This difference highlights that the rebinding of ligands from neighboring binding sites can influence measurements of intrinsic rates. Particularly, for weakly binding ligands, the lifetimes, thus the off rates, are overestimated for the systems in which binding sites are closer.

Overall, our MD simulations support the scaling analyses suggested in Section 2.1 for rebinding rates as well as total rebinding statistics. All regimes and their dependencies on two parameters, *h* and *s*, are in good agreement with the data extracted from our constant temperature simulations. Below we will discuss some implications of our results for various *in vivo* and *in vitro* situations.

## 3 DISCUSSION

Collective kinetic behavior of diffusing ligands can exhibit novel properties compared to that of a single ligand. In this study, we focus on the nonequilibrium rebinding kinetics of an ensemble of Brownian ligand particles in a confined volume that is initially free of ligands. Our study shows that nonsteady state on rates *k*_on_(*t*) and total number of revisits detected by each binding site 𝒩_coll_(*t*) depend on the two time scales imposed by the two intrinsic length scales of the corresponding system.

The first length scale is the largest spatial dimension of the diffusion volume. A steady-state kinetic behavior is reached only when the bulk density of diffusers becomes uniform in the corresponding volume. In experimentally typical flow cells, this length scale corresponds to the height of microchannel. For *in vivo* diffusion of signaling molecules throughout intercellular void, or in suspensions of cells or vesicles, this length scale can be related to average distance between receptor-bearing structures (i.e., 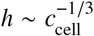, where concentration of cells is *c*_cell_). Once the steady state is reached, the on rate exhibits a time independent behavior as *k*_on_ ~ 1/*hs*^2^. The rebinding frequency in the steady state is characterized by an Einstein-Smoluchowski limit, which leads to a linearly increasing 𝒩_coll_ (Fig. 2).

The second length scale that we discuss in this work is the average separation distance between two binding sites, *s*, which is inversely proportional to the square root of grafting density of binding sites. At intermediate times (i.e., before the steady state is established), the on rate shows one or two power laws depending on *s*. For large values of *s*, due to the 3*d* escape process of ligands from their binding sites, the on rate exhibits a *k*_on_ ~ *t*^−3/2^ type of decay after the initial release of the ligand. Once the ligands diffuse to a distance larger than the separation distance *s*, above a threshold time of τ_s_, a quasi 1d diffusion process takes over with a smaller decay exponent of –½. For densely grafted binding sites (i.e., small *s*), the exponent 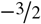 is completely smeared out, and the time dependence of the diffusion process is defined by a single exponent of –½ at intermediate times (Fig. 2a).

We also defined a time-dependent parameter 𝒩_coll_ (*t*) as the time integral of the on rate *k*_on_ (*t*) (more generally, the integral of raw dissociation data) to characterize rebinding kinetics. The parameter 𝒩_coll_ exhibits a novel plateau behavior on the scaling level at intermediate times for sparsely grafted binding sites (Fig. 2b). The plateau is a result of decreasing probability of finding any ligand near its binding sites during the 3*d* escape process. This behavior leads to a plateau behavior during which binding sites experience minimal number of collisions with the unbound ligands. Moreover, due to the integral form of Eq. 2, 𝒩_coll_ can be used to invoke the regimes in dissociation measurements otherwise difficult to observe due to relatively noisy statistics (see Figs. 4b and 5b).

The plateau expands to longer times if the binding sites are sparsely distributed since the terminal time for the plateau scales as τ_c_ ~ *s*^4^ (Eq. 4). The termination of the plateau regime is at τ_c_ instead of at τ_s_ ~ *s*^2^. We attribute this behavior to the non-uniform particle distribution near the surface at *t* < τ_c_; Only after particle the density becomes uniform near the reactive surface, binding sites can experience the incremental collision signals. This threshold time does not manifest itself in the dissociation data (Figs. 4a and 5a) and can be detected only from the cumulative consideration of the dissociation events ( (e.g., Figs. 4b and 5b)).

### 3.1 Relevance to experiments performed in microfluidic channels

Experimental studies exploring the kinetics of ligand-receptor interactions, or single-molecule-based biosensors, are commonly performed in microfluidic channels with well-defined dimensions. We will now discuss some consequences of our study on these experimental systems.

#### Low grafting density of receptors is essential to extract intrinsic kinetic rates in experiments

The measurable quantity in kinetic experiments such as SPR and fluorescence imaging, is the population of intact receptor-ligand complexes as a function of time, from which kinetic rates can be obtained. These apparatus cannot distinguish dissociation and subsequent association of ligands due to their finite-resolution windows. This means that, within the sampling time, a ligand-receptor pair can be broken and reform, possibly with new partners, and thus, contribute to the statistics as an intact complex. This can lead to artificially longer or shorter rates. Our study shows that if receptors are separated by small distances, the 3*d* escape process is quicker. Thus, rebinding of ligands desorbing from nearby receptors can alter intrinsic rates. We demonstrate this in our simulations (Fig. 8); Densely placed binding sites lead to longer lifetimes for ligands compared to the case in which binding sites are farther apart.

#### Association rates can have strong time dependence for weakly binding ligands

In the kinetic studies of receptor-ligand interactions (26) or in modeling signaling pathways (36), time- and concentration-dependent rates in master equations are common practices. Our study suggests that on rates can have non-trivial time dependence before the steady state is reached for diffusion-limited reactions and weakly binding ligands (e.g., a binding energy on the order of thermal energy). The time window within which this dependence continues is determined by the dimensions of experimental reservoirs, or average distance between ligand emitting and absorbing surfaces. As an example, a range of values around *h* = 10^2^ – 10^4^ *μ*m (4, 37, 38) leads to τ_*z*_ ≈ *h*^2^/*D* ≈ 10^2^ – 10^6^ s if we assume a diffusion coefficient of *D* = 100 *μ*m^2^/s for a ligand of size *a* = 1 nm (Table 1). The estimated values for τ_*z*_ are comparable to the lifetimes of molecular ligands (4), and the measurement taken earlier may not reflect true on rates, rather quantify an unrelaxed concentration quench. Note that, in the cases, τ_s_ > τ_*z*_, the regime with a ½ exponent in 𝒩_coll_ cannot be observed, and a direct transition to the long-time linear regime will be observed (Figs. 2 and 4b).

#### Separation distance brings about its own characteristic time scale

In single-molecule fluorescence imaging experiments of protein-DNA interactions, DNA binding sites are separated by distances on the orders of *s* ≈ 1 *μ*m (4, 6, 38). In SPR experiments, the distance between the surface-grafted receptors is often smaller and can be on the order of *s ≈* 10 nm (37, 39). Using, the same values for *D* and *a* as above, we can obtain some estimates as τ_s_ = 10^−6^ 10^−2^ s and τ_c_ = 10^−4^ 10^4^ s, respectively. While τ_s_, which characterizes the onset of the one-dimensional diffusion regime for on rate, is on the order of tens of miliseconds, τ_c_ can extend to hours since τ_c_ ~ *s*^4^ (Eq. 4). This wide spectrum of time scales suggests that with adequate design, receptor separations can be used to identify intermingled time scales in an heterogeneous system. For instance, biosensors can be prepared with multiple types of receptors (e.g., various nucleic acid sequences), each of which can have a distinct and tractable surface coverage level. Identification of signals coming from different sets of receptors can allow to interpret kinetic behavior of certain receptor-ligand pairs, if each separation distance distinctly manifests itself in dissociation kinetics.

#### Threshold timescales can be used to probe complex systems

The regimes that we discuss for experimentally measurable on rates and collision numbers can be used to extract average distance between receptors or receptor bearing surfaces. For instance, the threshold value τ_s_ in Eq. 5 can be utilized to obtain or confirm surface coverage levels of receptors without any prior knowledge if the decay of the dissociation data is not purely exponential. That is, as we discuss later, τ_s_ should be larger than the τ_off_.

#### Prospective experiments to untangle intermingled times scales

Recently, novel electrochemical-sensor applications based on the hybridization of a single stranded DNA binding site have been reported (15, 16). In these experiments, the voltage difference due to hybridization events of the grafted DNA by complementary strands in solution can be measured. Possibly, in these systems, extremely dilute binding site schemes can be constructed, thus, the time scales we discuss above can be validated experimentally. Indeed, as mentioned above, different nucleic acid strands can be grafted with varying separation distances, and in principle the resulting signals can be separated since our analysis suggest that each unique separation distance imposes its own terminal threshold times τ_s_ and τ_c_.

Another experiment setup that would be interesting could incorporate two SPR surfaces separated by a distance *h*. While one SPR surface can accommodate receptors saturated by ligands, opposing surface can have empty receptor sites, hence, create a “sink” for the ligands. Thus, both rebinding rates and arrival frequencies can be measured simultaneously. Signals on both surfaces could be compared by systematically varying the density of binding sites, surface separations, etc. Indeed, this or similar scenerios can be used to model diffusion of neutransmitters or growth factors *in vivo* (24**?**), since rebinding events on the secreting cells can become slower or faster depending on the number of receptors or their spatial distribution on target cell surfaces (13, 27, 40).

### 3.2 Signaling and communication via chemical gradients

The intercellular void formed by loosely packed cells can percolate to distances on the orders of microns (41, 42), which can lead to diffusion times on the orders of minutes. On the other hand, average (closest) distance between two neighboring cells can be on the orders of 10 nm (e.g., for synaptic cleft). Small molecules, such as cytokines, secreted from one cell can diffuse throughout these intercellular spaces and provide a chemical signaling system between surrounding cells. This type of communication is controlled by both secretion and transport rates (43). Indeed, recent studies suggest that spatiotemporal organizations of receptor and ligands can provide diverse signaling responses (44). In this regard, our result can be used to shed light on some aspects of chemical signaling processes *in vivo* as we will discuss next.

#### Time-dependent concentration near receptors can provide a feedback mechanism

Our results suggest that both on rates and total number of rebinding events are sensitive to time-dependent concentration fluctuations of ligands near secreting surfaces. According to our analyses, this time dependence ends when the ligands arrive opposing target surface (e.g., when neurotransmitters diffuse to the receptors of postsynaptic neuron). This suggests a feedback mechanism in which the secreting cells can determine the arrival of the released molecules to the target cells. This would be possible if the secreting cell bears receptors that are sensitive to the local concentration of the secreted molecule, possibly via time-dependent conformational (45, 46) or organizational (47–49) changes of membrane components. In this way, once the signal molecules reach their target surface, secreting cell can alter the signals depending on the rebinding regime experienced.

#### Exocytosis can be altered by concentrated vesicles or small openings

Our analysis shows that time-dependent on rates can reach their time-independent regimes faster, and the ligands return their initial position more often, if the ligands are closer to each other at the time of the initial release (see Figs. 2 and 5). In the process of exocytosis of small molecules, vesicles fuse with the plasma membrane and create an opening to release the molecules into intercellular space. One can imagine a scenario, in which, given the concentrations of contents of two vesicles are similar, a ligand released from the vesicle with a larger opening would return less to the original position (Fig. 2). If the vesicle opening is small, this would effectively lead to a smaller separation distance, thus, more return would occur. In fact, the amount of opening can also determine the efficiency of endocystosis (e.g., process of intake of ligand back to vesicle). In exocytosis, secreting vesicles can control the realease rates by changing modes of fusion (50). In accord with this concept, our calculations show that one order of magnitude decrease in the separation distance can increase the return rates by two orders of magnitude (Fig. 5a). Similarly, given the sizes of openings are roughly equivalent for two vesicles, more concentrated vesicle can lead to more collisions per unit time with the opposing cell surface, since the number of ligands per unit area is higher during the initial release for the concentrated vesicle (i.e., *k*_on_ ~ 1/*s*^2^). Similar arguments could be made to explain the observed differences in exocytosis rates induced by the fusion of multiple vesicles (51).

### 3.3 Finite residence time of ligands on binding sites

In the traditional view of molecular kinetics, the equilibrium constant of a bimolecular reaction (e.g., for a protein binding and unbinding its binding site along DNA molecule, or a drug targeting its receptor) is defined as the ratio of off rate *k*_off_ ≡ 1/τ_off_ and the corresponding on rate. As we discuss in Section 2.2.3, molecular ligands can have slow off rates (long residence times) that can intervene with the threshold times and regimes predicted by our scaling arguments. Moreover, these off rates can have strong concentration dependence (4, 5). Below, based on recent experimental findings (52), we will briefly discuss some implications of the finite residence times on our results.

#### Slow off rates can delay power laws

Due to various energetic and entropic components (7, 53), disassociation process of a ligand from its binding site can be considered as barrier crossing problem. This rare event manifests itself as a slower decay (compared to a diffusion-limited case) in dissociation curves, which is usually fited by either exponential or nonexponential curves (30) to extract dissociation rates. For ligands with strong affinity towards their binding sites (i.e., τ_off_ /τ_s_ ≫ 1), this slow decay can occlude the power laws that we discuss, depending on the residence time. In Fig. 9, we demonstrate the possible effects of the residence times on our calculations with an illustrative diagram in the dimensions of *s*/*h* and τ_off_/τ_s_.

In Fig. 9, if the residence time of a ligand is short compared the inter-site diffusion time (i.e., τ_off_/τ_s_ < 1), the ratio of *s*/*h* determines which power law or laws can be dominant at intermediate times. For instance, for *s*/*h* < 1, which is the common scenario in SPR experiments, both of the decay exponents can be apparent. In other single-molecule experiments, for which 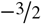 type of exponent can be observable as long as there are enough empty sites for dissociating ligands (i.e., τ_s_ > τ_off_). If the residence and inter-site diffusion times are comparable (i.e., τ_off_ **/**τ_s_ ≈ 1), the regime of *k*_on_ ~ *t*^−3/2^ is not observable, and the on rate decays by –½ exponent until the steady-state regime. For a dense array of binding sites, a non-exponential decay can also arise as a result of correlated rebinding events (22).

#### Time-dependent concentration can induce time-dependent facilitated dissociation

The recent studies of protein-DNA interactions have shown that off rates, 1/τ_off_, have a strong dependence on concentration of unbound (free) proteins in solution (4, 6, 8). According to this picture, free ligands in solution can accelarate dissociation of bound proteins by destabilizing the protein-DNA complex (4, 54). Our study shows that, for a concentration quench, concentration of ligands near the binding site changes with time prior to steady state. The time-dependent concentration can lead to time-dependent facilitated dissociation and shorten the lifetimes of ligands on their binding sites in a time-dependent manner, particularly in the reaction limited scenario. This scenario can be more important when binding sites are too close to each other, since cross rebinding events can induce more facilitated dissociation and further shorten the residence times of the ligands. We do not expect to see this mechanism in our simulations since the facilitated dissociation mechanism usually requires ligands with multivalent nature, which can exhibit partially bound state (54). Future studies can address this mechanism by considering, for instance, dimeric ligands.

## AUTHOR CONTRIBUTIONS

All authors contributed equally to this work.

## ACKNOWLEDGMENTS

A.E. acknowledges Edward J. Banigan and Ozan S. Sarıyer for their careful readings of the manuscript, and Reza Vafabakhsh for bringing important literature on synaptic release to our attention. We acknowledge The Fairchild Foundation for computational support. J.F.M. acknowledges the NIH grants CA193419 and U54DK107980, and M.O.d.l.C. acknowledges the NSF grant DMR 1611076.

## A. Derivation of on rates via Gaussian distribution

Here we derive expression for *k*_on_ and 𝒩_coll_ by using a Gaussian spatial distribution for ligands. Consider at time *t* > 0, the probability distribution for a set of identical particles in *d* dimensions evolves from a dirac delta distribution at the origin as

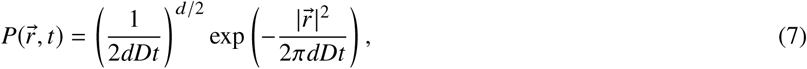

where 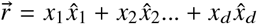 is the position vector in *d* dimensions. The weight of the distributions in Eq. 7 at position 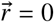 provides a probability for diffusing particles to revisit the origin

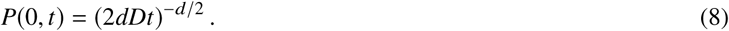

At time *t*, the total number of the revisits can be obtained by integrating Eq. 8

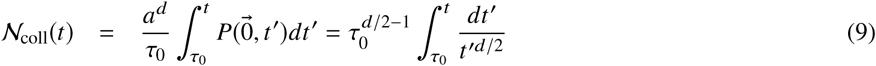

According to Eq. 9, the returning probability, 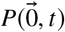, can also be considered as the rate of revisits, *k_on_*, at the origin 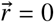 at a given period *T*.

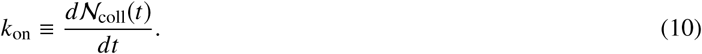

After dimensional adjustment, Eq. 10 can also be written as

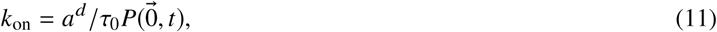

From Eq. 11, the on-rates are

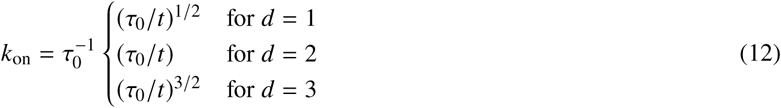

If the integral in Eq. 9 can be performed to obtain the expressions for 𝒩_coll_(*t*) as

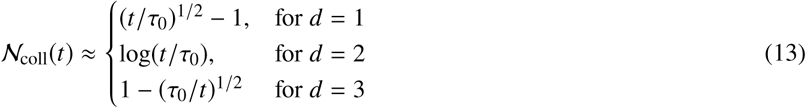

Using Eqs. 12 and 13, the scaling arguments presented in the main text can be obtained for each step of the diffusion process. Note that even though a 2*d* scenario has been realized for the current problem, it has been show before that diffusion profile of protein particles that are initially positioned along a one dimensional chain obeys a logarithmic revisit rates (10).

## APPENDICES

### B. MD simulation details

In the simulations, *n*_0_ = 400 – 6400 binding sites separated by a distance *s* are placed on a planar surface composed of square lattice of beads of size *a*. To model ligands, *n*_0_ beads of size 1σ ≈ *a* are placed at contact with binding sites, where σ is the unit distance in the simulations.

The steric interactions between all beads are modeled by a truncated and shifted Lennard-Jones (LJ) potential, also known as WCA,

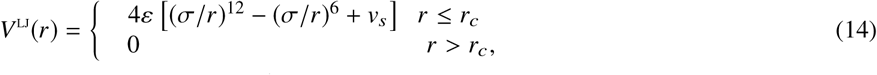

where *r_c_* is the cutoff distance. A cut-off distances of *r_c_* /σ = 2^1/6^, 2.5 are used with a shift factor *v_s_* = 1/4 for the interactions between all beads unless otherwise noted. The interaction strength is set to *ε* = 1*k*_B_*T* for all beads, where *k*_B_ is the Boltzmann constant, and *T* is the absolute room temperature. For attractive cases, the cut-off distance is set to *r_c_*/σ = 2.5 and the strength of the potential is varied between *ε* = 0.5 – 3*k*_B_*T*

All MD simulations are run with LAMMPS MD package (33) at constant volume *V* and reduced temperature *T_r_* = 1.0. Each system is simulated for 10^6^ to 2 × 10^9^ MD steps. The simulations are run with a time step of ∆*t* = 0.005τ, where the unit time scale in the simulations is τ ≈ τ_0_. The data sampling is performed by recording each 1, 10, 10^2^, 10^3^ and 10^4^ steps for MD intervals 0 – 10^2^, 10^2^ – 10^3^, 10^3^ – 10^4^, 10^4^ – 10^5^ and 10^5^ – 10^8^, respectively. The monomeric LJ mass is *m* = 1 for all beads. The temperature is kept constant by a Langevin thermostat with a thermostat coefficient *γ* = 1.0τ^−1^.

The volume of the total simulation box is set to *n*_0_(*s*^2^ *h*)σ^3^, where the vertical height is *h/*σ = 12.5 – 2000. Periodic boundary conditions are used in the lateral (*x*̂ and *y*̂) directions and at *z* = *h* simulation box is confined by a surface identical to that at *z* = 0. VMD is used for the visualizations (55).

In the fitting procedures, a weight function inversely proportional to the square of the data point is used. Error bars are not shown if they are smaller than the size of the corresponding data point.

## SUPPLEMENTARY MATERIAL

An online supplement to this article can be found by visiting BJ Online at http://www.biophysj.org.

